# Ultra-Low-Cost Integrated Silicon-based Transducer for On-Site, Genetic Detection of Pathogens

**DOI:** 10.1101/2020.03.23.002931

**Authors:** Estefania Nunez-Bajo, Michael Kasimatis, Yasin Cotur, Tarek Asfour, Alex Collins, Ugur Tanriverdi, Max Grell, Matti Kaisti, Guglielmo Senesi, Karen Stevenson, Firat Güder

## Abstract

Rapid screening and low-cost diagnosis play a crucial role in choosing the correct course of intervention *e.g.,* drug therapy, quarantine, no action etc. when dealing with highly infectious pathogens. This is especially important if the disease-causing agent has no effective treatment, such as the novel coronavirus SARS-CoV-2 (the pathogen causing COVID-19), and shows no or similar symptoms to other common infections. We report a silicon-based integrated Point-of-Need (PoN) transducer (TriSilix) that can chemically-amplify and detect pathogen-specific sequences of nucleic acids (NA) quantitatively in real-time. Unlike other silicon-based technologies, TriSilix can be produced at wafer-scale in a standard laboratory; we have developed a series of methodologies based on metal-assisted chemical (wet) etching, electroplating, thermal bonding and laser-cutting to enable a cleanroom-free low-cost fabrication that does not require processing in an advanced semiconductor foundry. TriSilix is, therefore, resilient to disruptions in the global supply chain as the devices can be produced anywhere in the world. To create an ultra-low-cost device, the architecture proposed exploits the intrinsic properties of silicon and integrates three modes of operation in a single chip: i) electrical (Joule) heater, ii) temperature sensor (*i.e.* thermistor) with a negative temperature coefficient that can provide the precise temperature of the sample solution during reaction and iii) electrochemical sensor for detecting target NA. Using TriSilix, the sample solution can be maintained at a single, specific temperature (needed for isothermal amplification of NA such as Recombinase Polymerase Amplification (RPA) or cycled between different temperatures (with a precision of ±1.3°C) for Polymerase Chain Reaction (PCR) while the exact concentration of amplicons is measured quantitatively and in real-time electrochemically. A single 4-inch Si wafer yields 37 TriSilix chips of 10×10×0.65 mm in size and can be produced in 7 hours, costing ~US $0.35 per device. The system is operated digitally, portable and low power – capable of running up to 35 tests with a 4000 mAh battery (a typical battery capacity of a modern smartphone). We were able to quantitatively detect a 563-bp fragment (Insertion Sequence IS*900*) of the genomic DNA of *M. avium* subsp. *paratuberculosis* (extracted from cultured field samples) through PCR in real-time with a Limit-of-Detection of 20 fg, equivalent to a single bacterium, at the 30^th^ cycle. Using TriSilix, we also detected the cDNA from SARS-CoV-2 (1 pg), through PCR, with high specificity against SARS-CoV (2003).

## 1. Introduction

Despite the advancement of diagnostic technologies targeting nucleic acids (NA), there are still no rapid, handheld, low-cost, easy-to-use and integrated solutions for the testing of infectious diseases at the Point-of-Need (PoN). This is unfortunately the case for pathogens infecting humans, animals or even plants. This large gap in the diagnostic workflow hampers attempts to contain infectious pathogens from spreading by rapid and early detection. This technological gap has become, once again, evident with the spread of COVID-19, the diagnosis of which continues to depend heavily on centralized laboratories with specialized personnel and facilities, which in turn slows down testing and delays treatment.

NA make excellent targets for the direct detection of pathogens due to their high specificity. Unlike other biomarkers, such as antibodies or non-genetic molecules originating from pathogens, NA can be chemically amplified, enabling direct detection of low numbers of pathogens (down to single organisms). Hence tests targeting NA tend to be exceptionally sensitive. NA-based reagents (*e.g.,* primers) for NA testing can also be produced synthetically rapidly and on a large scale; DNA-based molecules, in particular, are highly stable, therefore, do not require a cold-chain for storage, making it especially suitable for PoN testing.^1^ Ability to produce NA quickly also increases the speed of development and deployment of new test kits to health systems, providing much needed diagnostic capabilities for the detection of novel pathogens (such as SARS-CoV-2). Despite the massive advantages, on-site testing of NA has been limited.

There are several molecular methods to amplify NA for detection with high specificity *e.g.,* polymerase chain reaction (PCR), loop-mediated isothermal amplification (LAMP), strand-displacement amplification (SDA), recombinase-polymerase amplification (RPA). While PCR requires thermocycling; emerging isothermal methods such as LAMP and RPA, do not, removing the need for sophisticated instruments.^2,3^ On the other hand, PCR is a simple process, requiring few reagents. It is also the gold standard for NA-based laboratory diagnostics with a solid support infrastructure. Amplification strategies are traditionally combined with fluorescence-based optical methods of detection to produce a quantitative analytical signal. Optical methods, even though sensitive, require expensive instruments.^4^ The fluorescent labels used in optical detection are also difficult to handle and susceptible to photobleaching. Optical detection systems have been difficult to miniaturize and largely limited to centralized laboratories, however, some early concepts have been proposed by various groups in the literature.^5–10^

Despite low-cost onsite testing for infectious diseases is being the holy grail of NA diagnostics, there are still no inexpensive and handheld solutions in the market that can provide truly portable, rapid NA detection. Commercially available, benchtop optical lab instruments such as GeneXpert (Cephied) have already made a substantial difference in the speed of NA-based diagnosis of infectious diseases but the high cost of the instrument/tests, large size and power consumption have prevented the adoption of these systems for use in the field.

The ideal, miniaturized, low-cost portable detection system for NA must be able to heat up the sample from the patient to the desired temperature setpoint with high precision (using a heater and temperature sensor) and measure the results of the amplification reactions quantitatively, all in an integrated fashion. Using cleanroom-based semiconductor fabrication methods, such systems have been reported;^11–15^ however, these devices require advanced methods of microfabrication that can only be performed in a cleanroom, such as photolithography, vacuum etching/deposition etc. They also do not exploit the intrinsic properties of the semiconductor materials that are used as substrates (*i.e.* Si itself can be used both as a temperature sensor and electrical heater). Hence the devices reported are complex and expensive. Because semiconductor foundries are mainly located in East Asia, manufacturing and logistics are susceptible to disruptions in the supply chain due to unexpected events (*e.g.,* COVID-19 pandemic). There is, therefore, still no viable low-cost, rapid, quantitative, integrated handheld NA testing solution available for use in the field with the potential for market wide availability.

In this work, we report an integrated silicon-based Point-of-Need tri-modal NA transducer (TriSilix) that can chemically-amplify and detect pathogen-specific sequences of NA quantitatively in real-time (**Figure 1** and Supporting Information **Video V1**). Unlike other silicon-based technologies, TriSilix can be produced at wafer-scale in a standard laboratory and does not require fabrication in an advanced semiconductor foundry. Hence it is resilient to disruptions in the global supply chain as the devices can be produced anywhere in the world. To achieve cleanroom-free, low-cost fabrication, we have developed a series of fabrication methodologies based on wet etching to form porous Silicon, electroplating, thermal bonding and laser-cutting (Figure 1A). To reduce costs and complexity, TriSilix exploits the intrinsic properties of the semiconductor Si which can be used as a resistive heating device and thermistor simultaneously. TriSilix has three modes of operation: i) electrical (Joule) heater, ii) thermistor with a negative temperature coefficient that can provide the precise temperature of the sample solution during reaction and iii) label-free electrochemical sensor for detecting target NA with methylene blue as a redox-active reporter. TriSilix works with both cyclic and isothermal methods of amplification. We demonstrate that TriSilix can detect NA from both bacteria (*Mycobacterium avium* subspecies *paratuberculosis*) and viruses (SARS-CoV-2) with high specificity.

**Figure 1.**
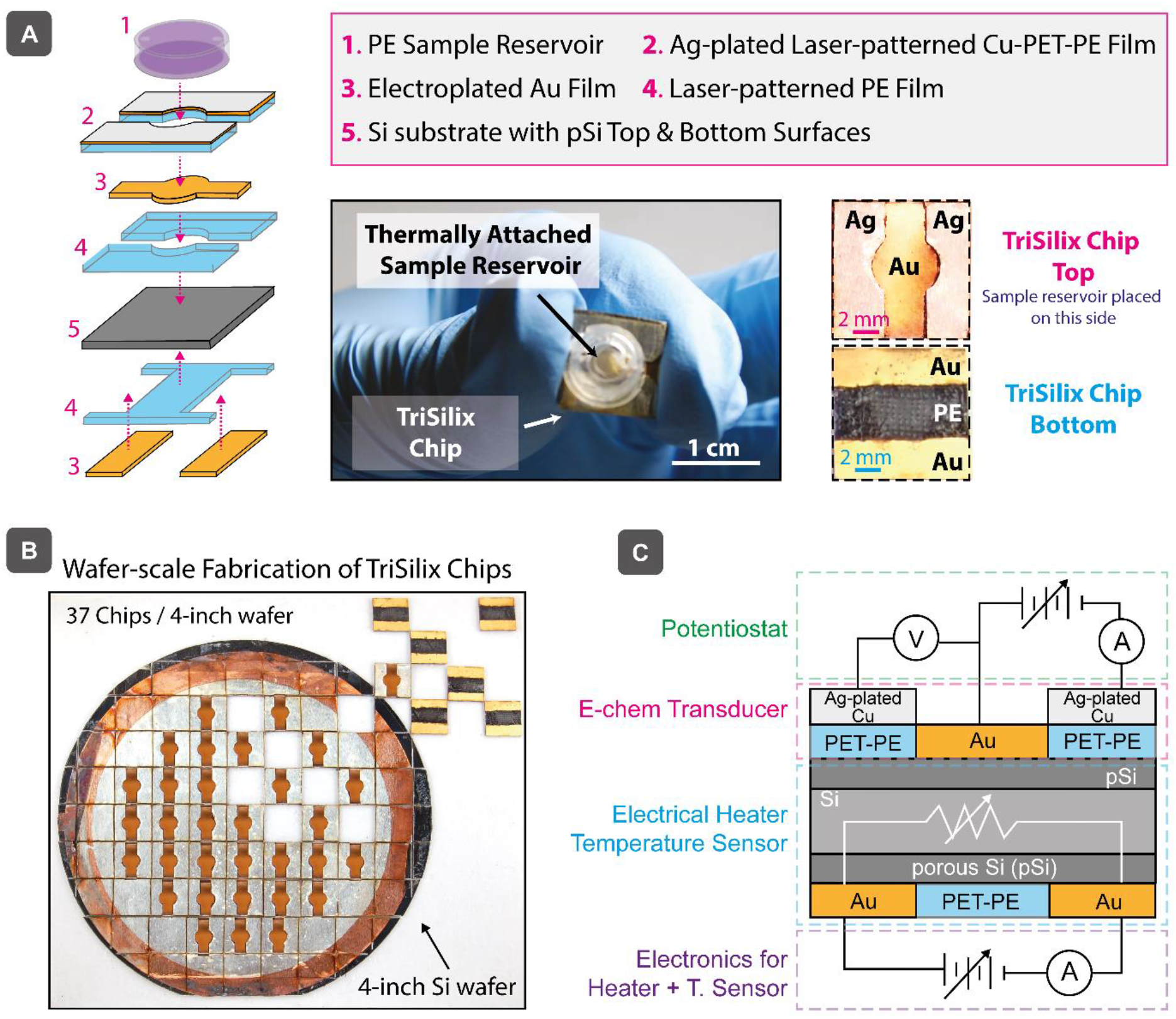
**(A)** Schematic illustration of construction of a TriSilix chip and photographs of the actual device. **(B)** Wafer-scale fabrication of TriSilix using a 4-inch Si wafer. Each wafer yield 37 chips. (**C)** Schematic illustration of the functional building blocks of TriSilix that provide trimodal operation for integrated nucleic acid amplification and detection.

## 2. Fabrication of TriSilix Chips

The fabrication of multilayered, composite TriSilix chips (dimension of each chip: 10×10×0.65 mm) starts with a Si wafer.^16^ In this study we used lightly-doped p-type 4” Si wafers, however, wafers with a larger diameter would also work.

First, each wafer is plated electrolessly for 20s (in 80 μM KAuCl_4_ and 0.5% HF) to form a thin layer of gold particulate film. The wafer is then placed inside an etching bath containing an aqueous solution of H_2_O_2_ and HF with a ratio of 1:20 v/v (30% H_2_O_2_: 10% HF) to perform metal-assisted chemical etching (MACE) of Si for 10 min.^17–19^ This process forms a 500 nm thick nanoporous Silicon (pSi) layer on each side of the wafer (**Figure 2A**). pSi plays a critical role in the fabrication of TriSilix. The porous surface allows electroplating of high-quality metal films on the surface of the Si substrate by creating an interlocking, high porosity surface to improve adhesion **(Figures 2B & C)**. Without this step, the metal films electroplated do not adhere to the surface of the substrate.^19^ The pSi layer also allows thermal bonding of sheets of polymer films in an ordinary heat press after patterning through laser-cutting (**Figure 2D**). After the formation of pSi surface, two layers of polyethylene terephthalate-polyethylene (PET-PE) are thermally bonded on the bottom and top surfaces of the Si wafer by heat pressing at 180 °C for 5 min. The heat pressed layers define the shape of the Working Electrode (WE) on the top surface and electrical contacts for Joule heating/resistance measurements on the bottom surface of the wafer. The patterned, and heat pressed polymer sheets essentially act as masking layers for electroplating of the metal electrodes on the pSi surface equivalent to photolithography-based patterning in conventional microfabrication. Next, the unmasked areas of the pSi surface were cleaned in 5% HF and electroplated by Au in a bath containing an aqueous solution of 10 mM KAu(CN)_2_/KCN for 10 min under a constant current of 10 mA versus Pt electrode (5×5 cm) yielding ~100 nm thick porous Au films. To achieve high uniformity when electroplating across the wafer, we have designed, and 3D printed a custom, electroplating platform that allows circular electrodes to be attached around the perimeter of the Si wafer (**Figure S1**). To form a three-electrode electrochemical cell on the top surface, the Counter (CE) and Reference Electrodes (RE) were created by heat pressing an Ag-plated Cu-PET-PE film, patterned once again by laser cutting. Ag plating was performed in an aqueous solution of 10 mM AgCN/KCN under an applied current of 20 mA versus Pt electrode (using the electroplating holder). Once the basic device structure was created, the wafer was laser-diced into individual chips (37 chips / 4” Si wafer, Figure 1B). Using the Randles–Sevcik equation with experimentally measured gradient values of 0.71 ± 0.08, we determined that the Au film has a 2x larger electroactive area than the geometrically defined area due to the high porosity of the Au film, which is beneficial for high-performance electrochemical analysis (**Figure S2**). A sample reservoir (polyethylene; diameter: 5 mm; volume: 40 μl, Figure 1A) with two holes was thermally bonded across the three-electrode electrochemical cell at 110±10 °C. The sample reservoir is specifically designed to prevent the evaporation of the solvent during amplification of DNA at elevated temperatures. The final TriSilix chip has a thickness of 490 μm (N=7) in the WE region and 650 μm in the RE/CE region (N=7) without the sample reservoir. Figure 1C shows a schematic illustration of the functional building blocks of TriSilix that provide trimodal operation: The three-electrode electrochemical cell fabricated on the top surface is connected to an electrochemical reader (potentiostat) to provide an electrochemical signal concerning the detection of NA (Mode 1). The electrodes fabricated at the bottom of the device are connected to custom electronics to enable heating (Mode 2) and temperature sensing (Mode 3) for thermoregulation.

**Figure 2.**
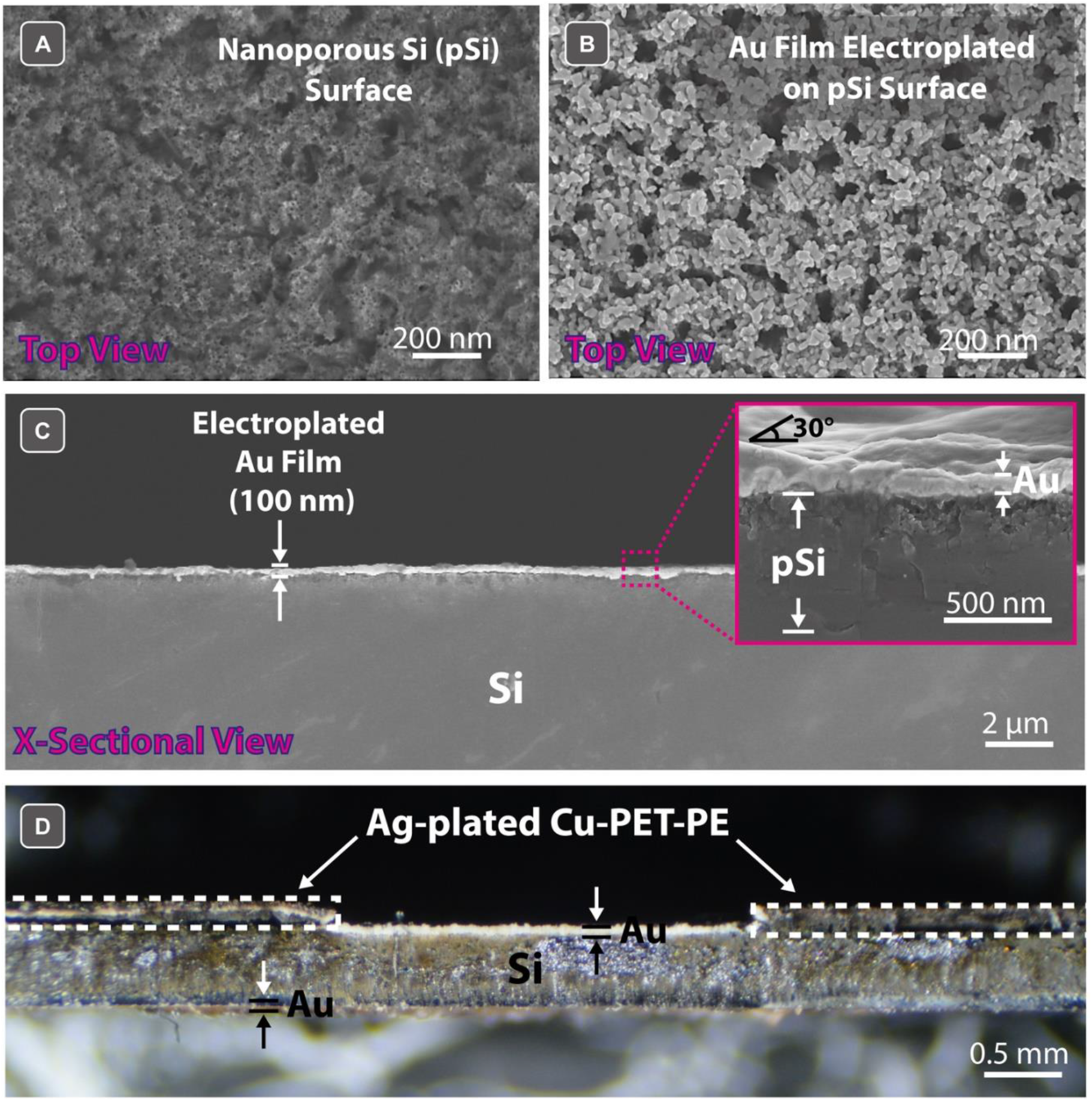
Scanning electron micrographs of the surface of **(A)** Si wafer after metal-assisted chemical etching with Au catalyst producing a nanoporous Si surface (pSi). (**B)** pSi surface after electroplating with gold. **(C)** Cross-section of the sample shown in (B). The inset is a close-up view of the cross-section where the Au and porous Si layers are clearly visible. **(D)** Optical micrograph of the cross section of a TriSilix Chip with thermally bonded sheets of polymer films. Note that while the cross-sectional image shown in (C) was cleaved, (D) was laser-cut hence the roughness.

## 3. Characterization of Temperature Transduction

NA amplification reactions require maintaining the sample at a temperature setpoint with high °precision. For PCR, the duration of each heating step must also be carefully controlled. Because the TriSilix uses Si, the substrate itself can be used both as an electrical heater and temperature sensor (*i.e.* thermistor) without adding any additional components for temperature transduction. Si, a semiconductor, heats up when an electrical current passes through it (known as Joule heating). Because Si has a high thermal conductivity (~150 W m^−1^ °K^−1^ at 300 °K), the substrate can be heated uniformly regardless of the path the current flows. The electrical resistance of Si also varies with temperature with a negative correlation (within our experimental range of temperatures from RT to 110 °C); the electrical resistance of Si drops with increasing temperature due to generation of mobile charge carriers allowing the use of Si substrate itself as a sensitive sensor of temperature. ^20–22^

We have applied electrical currents in the range 0-400 mA between two Au electrodes deposited on the bottom of TriSilix chip to heat up the device electrically (the experiments were performed at room temperature; ~25 °C). During this experiment, we used a thermal camera (FLIR E4) to measure the temperature across the chip as a reference measurement. As illustrated in **Figure 3A,** TriSilix chip can be heated up to 110 °C, electrically. The relationship between the current applied and substrate temperature was linear (with a positive slope) at steady-state with high repeatability (slope: 232.3±8.7 °C A^−1^; R^2^ = 0.9971, N=5).

**Figure 3.**
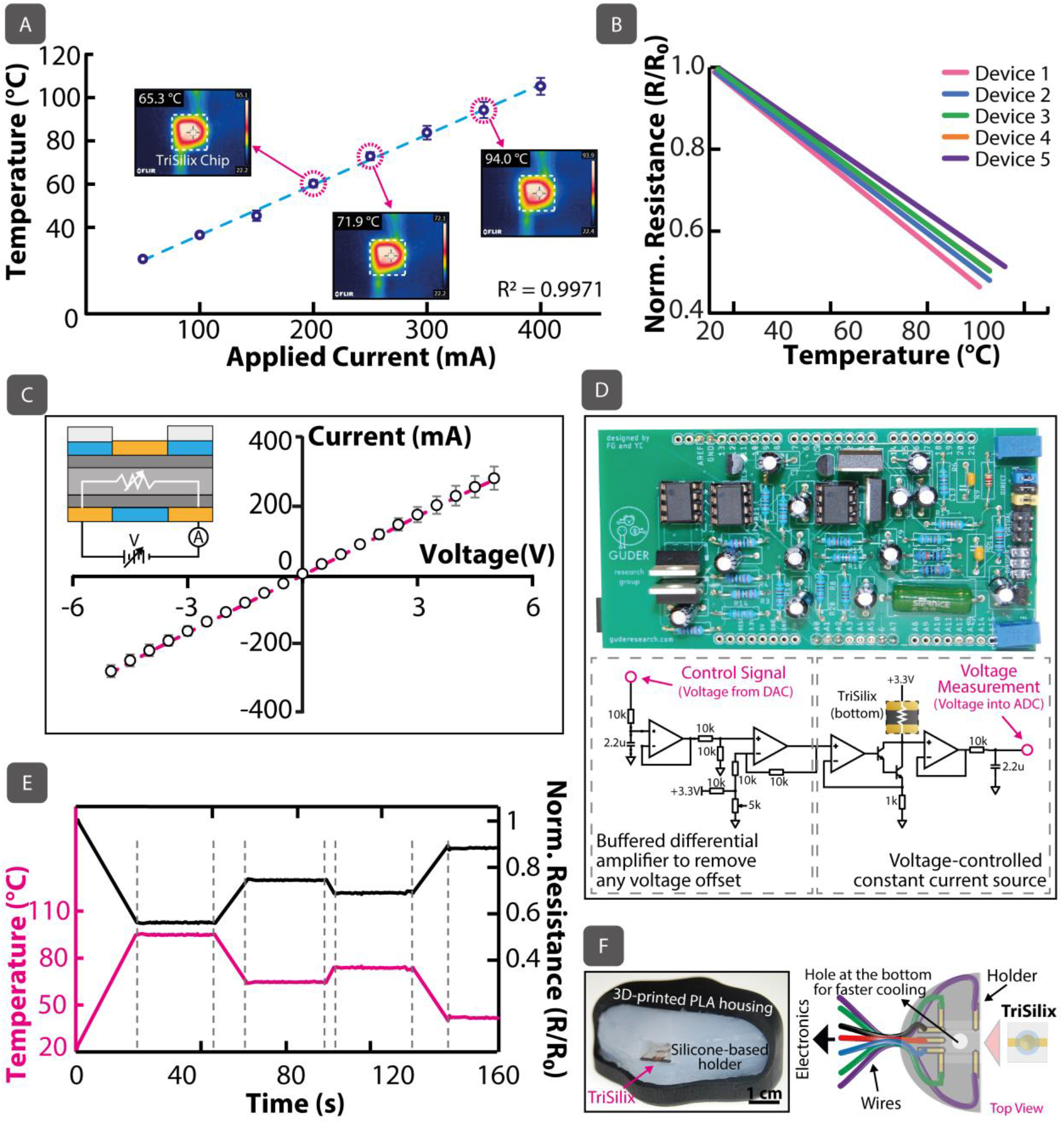
**(A)** Relationship between temperature and (constant) current applied when TriSilix is used as an electrical heater (N=5). The temperature measurements were cross correlated with a thermal camera as a reference measurement. **(B)** Temperature vs normalized resistance for five TriSilix chips. The relationship between the normalized resistance (resistance/initial resistance) were linear for all devices with a negative slope. **(C)** I-V measurements across the Au-Si-Au junction which yielded a linear, Ohmic relationship between −5V and 5V (R^2^=0.9996, N=5). **(D)** Photograph (top) and simplified circuit schematic diagram (bottom) of the custom electronics used for heating and temperature sensing. **E)** Temperature program to maintain the TriSilix chip at a temperature setpoint over time using software control and custom electronics. This program is an exemplar fifth cycle of a TriSilix PCR program in which the sample is kept at 30 sec for each step at 94 °C, 63 °C and 72 °C before being lowered to 40 °C for the electrochemical measurement. **F)** Photograph (left) and schematic illustration (right) of the silicone-based holder with embedded contacts to run TriSilix in trimodal operation. Error bars=SD.

At room temperature, the electrical resistance of a batch of TriSilix chips was 5.4±0.6 Ω (N=37) which varied linearly (with a negative slope of −6.1±0.4×10^−3^ °C^−1^; R^2^ = 0.9991, N=5) with the temperature measured within the range from room temperature to 110 °C (**Figure 3B**). By measuring the electrical resistance of the Si substrate, using the two electrical contacts at the bottom of the chip, the temperature of the TriSilix chip can be precisely identified after calibration against a reference measurement (see Supporting Information **Section S3** for more information on the calibration of the TriSilix thermistor). Temperature sensing is important for correct and fast execution of a NA amplification program for at least two reasons: i) Temperature can be maintained at a precise setpoint using a control algorithm such as proportional–integral–derivative control loop. ii) When cooling (passively) or heating (actively), the speed of reaching a temperature setpoint can be maximized as both heating and cooling depend on the outside temperature and device packaging (*e.g.,* the construction of the holder or cartridge). By only relying on the power applied, temperature control cannot be achieved with high precision.

Although we expected formation of Schottky barriers and non-linear electrical characteristics across the Au-Si interface, the I-V measurements (**Figure 3C**) demonstrated that the junction exhibits relatively ohmic behavior with a linear I-V relationship (R^2^=0.9996) between −5V and +5V. The Si substrate plus the Au contacts can, therefore, be modeled as a simple variable resistor, the magnitude of which changes with temperature. We have designed a custom electrical circuit (**Figure 3D**) and MATLAB-based graphical user interface (**Figure S4**) to control the temperature of the TriSilix chip at a precise setpoint through software control. The circuit (Figure 3D) implements a voltage controlled constant current source which can be adjusted digitally using a low-cost microcontroller (Arduino Due). This circuit can also measure the voltage drop across the TriSilix chip; when a constant current is applied, the resistance (hence the temperature) of the TriSilix thermistor can be calculated using the voltage reading, and Ohm’s law. Using the MATLAB program and the custom electrical circuit board, we were able to run a temperature program (**Figure 3E**) and maintain the TriSilix chip at a given temperature setpoint over time. This is crucial for both cyclic and isothermal NA amplification reactions.

## 4. Characterization of Electrochemical Sensing

We produced a thermally-stable silicone-based holder encased in a 3D printed polylactic acid (PLA) housing (**Figures 3F and S5**) to characterize the performance of electrochemical sensing using the TriSilix chip. The silicone holder also contained five embedded gold-plated stainless-steel electrodes to make electrical contacts with the three electrodes positioned at the top for electrochemical NA sensing and four at the bottom for heating and temperature sensing.

First, we characterized the electrochemical redox processes for methylene blue (MB) using cyclic voltammetry (**Figure 4A**). We chose MB because the electrochemical approach we used for the detection of DNA involves the use of MB^23–25^ as an intercalating redox reporter. During NA amplification, MB is intercalated between guanine-cytosine base pairs of the double-stranded DNA (ds-DNA) which provides an electroanalytical signal correlated with the concentration of ds-DNA in the sample. We prepared a 125 μg mL^−1^ solution of MB in 10 mM phosphate-buffered saline pH 7 (PBS) and swept across a range of potentials between −400 to 200 mV at a scan rate of 100 mV s^−1^ to characterize the electrochemical processes involving MB and electrodes. We determined that the anodic peak current, originating from the oxidation of MB on the electrode surface, appears at −67±2 and cathodic peak (due to reduction) at −97±4 mV versus Ag (Figure 4A). The absence of peaks with nearly overlapping anodic and cathodic curves in buffer alone indicates that the electrodes are electrochemically and mechanically stable and high performance (*e.g.,* low capacitive current). We have performed (**Figure 4B**) square wave voltammetry (SWV), a substantially more sensitive electroanalytical method, to measure the concentration of MB in PBS in a range between 0 - 125 μg mL^−1^ using a potential window from −500 mV to −250 mV versus Ag (corresponding to the anodic process). The SWV measurements using TriSilix produces an electroanalytical signal (peak current intensity) that is linearly related to the concentration of MB in the range 0.5 μg mL^−1^ – 80 μg mL^−1^ with an R^2^ = 0.9963 (**Figure 4C** – curve denoted by ‘RT’). We have also characterized the analytical performance of TriSilix for performing SWV measurements when the chip was operated at elevated temperatures. To prevent evaporation of the solvent (*i.e.* water) from the sample solution, a small amount (10 μL) of mineral oil was added to the reservoir. Since both isothermal and cyclic NA amplification reactions require heating; the effect of temperature on the electrochemical measurements is important. As shown in Figure 4C, the peak current intensity (measured from the recorded SWVs) increased two to four times in comparison to room temperature when operated at higher temperatures; this is due to enhanced transport of the analyte to the surface of the electrode.^26–28^ With increasing temperature, however, the error bars also widen, indicating that the precision of the measurement decreases. We speculate that this could be related to thermal and electrical crosstalk between the heater and the electrochemical sensing structures which introduce additional noise to the electroanalytical measurements at higher temperatures. We have also studied the thermal stability of MB over time (**Figure 4D**). In this experiment, we added MB to the PCR and RPA mastermix solutions to simulate the conditions during isothermal and cyclic amplification reactions. As illustrated in Figure 4D (and **Figure S6**), MB remained relatively stable both at 40 °C over 45 mins (*i.e.* RPA conditions) and 35 cycles of PCR conditions (30 sec for each step at 94 °C, 63 °C and 72 °C; every 5^th^ cycle, the temperature is reduced to 40 °C for the measurement to increase precision as illustrated in **Figure S7**) indicated by the small spread of the measurements. The SWV measurements at elevated temperatures produced similar results to the measurements taken at room temperature (25°C) in terms of repeatability, in agreement with the findings in the literature by other groups although we did observe a slight decrease in stability for the PCR conditions (*i.e.* higher temperatures).^23^ The results of the thermal stability experiments indicate that MB is a sufficiently stable redox-active reporter for use in NA amplification reactions at elevated temperatures.

**Figure 4.**
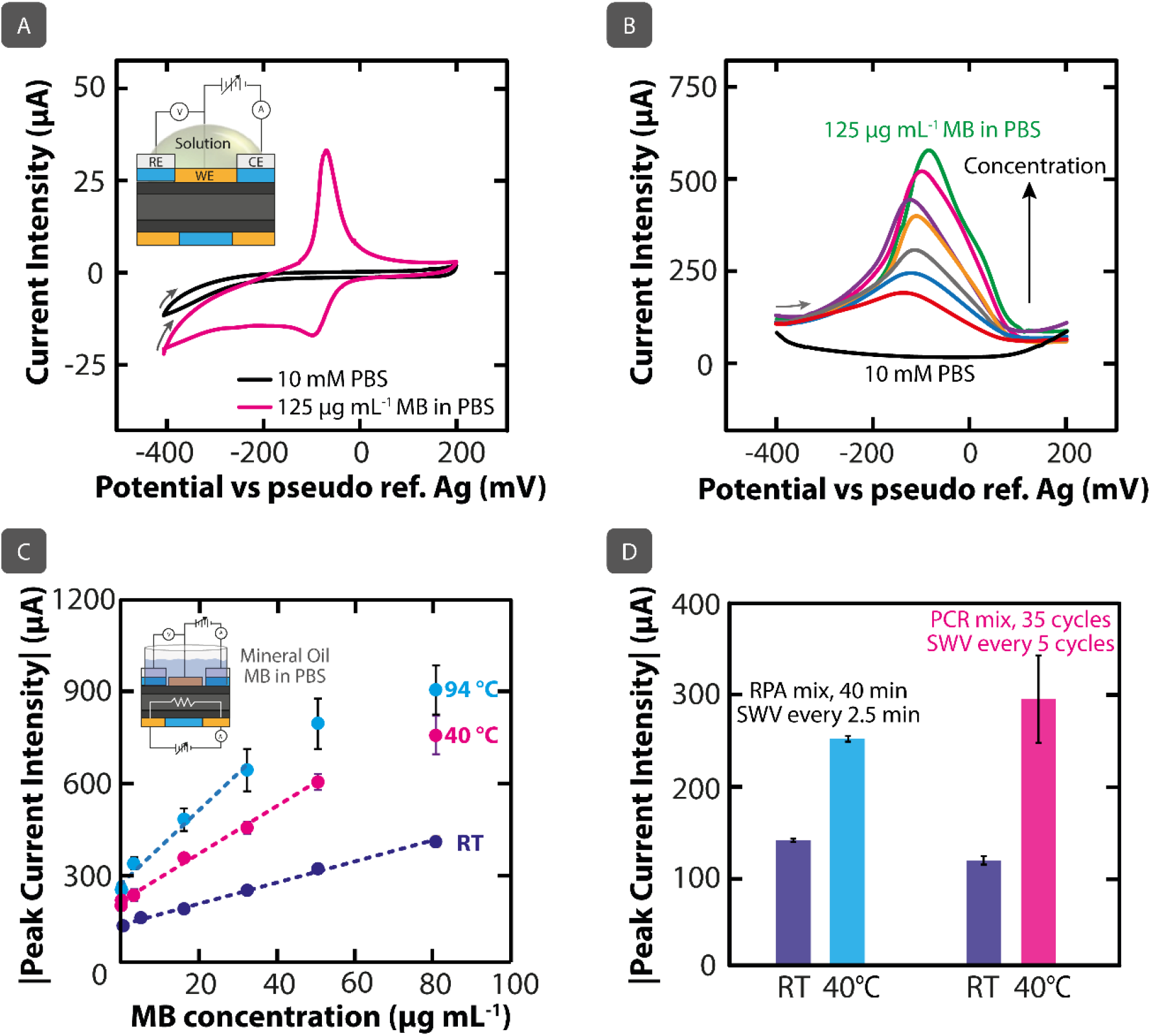
**(A)** CVs recorded in 0 and 125 μg mL^−1^ aqueous solutions of MB in PBS. Scan rate=100 mV s^−1^. (**B)** SWVs recorded in MB solutions in PBS at room temperature in a range of concentrations from 0 to 125 μg mL^−1^. (**C)** Calibration curves of peak current intensities vs concentration from SWVs recorded in MB solutions in PBS in a range of concentrations from 0 to 80 μg mL^−1^ at different temperatures. Error bars=SD, N=5. **(D)** Histogram showing average values from SWVs recorded in RPA mastermix and PCR mastermix at room temperature and under NA amplification conditions. Error bars correspond to standard deviations from peak current intensities measured from 17 (RPA) and 8 (PCR) SWVs.

## 5. Real-time Quantitative Detection of DNA During Amplification Reactions

Using the TriSilix chip, we have performed real-time and quantitative RPA (isothermal, qRPA) and PCR (cyclic, qPCR) amplification of DNA using its temperature transduction capabilities and measured the concentration of reaction products – smaller fragments of ds-DNA, *i.e.* amplicons – using the electrochemical DNA sensor, quantitatively and in real-time. During amplification, the number of amplicons increases exponentially with time if the DNA target is present in the sample. When new amplicons are generated, MB interacts with the G-C pairs and can no longer participate in the electron-transfer reactions with the working electrode hence the electroanalytical signal, originating from the redox-active reporter, decreases with increasing concentration of amplicons (and the target DNA in the original sample).

We performed qRPA using TriSilix with MB as the redox-active reporter. We used the TwistAmp Basic kit (TwistDx, UK) with the positive (3.2 kbp template, 144-bp amplicons) and negative (100-bp CTX-M ESBL; extended-spectrum beta-lactamases) DNA controls in the experiments. We have slightly modified the mastermix solution provided in the kit to contain 10 μg mL^−1^ of MB to enable electrochemical detection in real time. 30 μL of the modified-mastermix was introduced into sample reservoir of TriSilix followed by the addition of 10 μL of mineral oil to prevent evaporation. The sample was heated to 40 °C for 40 min and a square-wave voltammogram was recorded every 2.5 min during the reaction. As shown in **Figure 5A,** while the electroanalytical signal (normalized peak current intensity) remained relatively unchanged for the negative control, the positive control (50 – 150 nM according to the manufacturer) showed a large drop in signal in the form of a characteristic sigmoidal profile. This result demonstrates that TriSilix can perform real-time and quantitative DNA analysis through (isothermal) RPA reactions.

**Figure 5.**
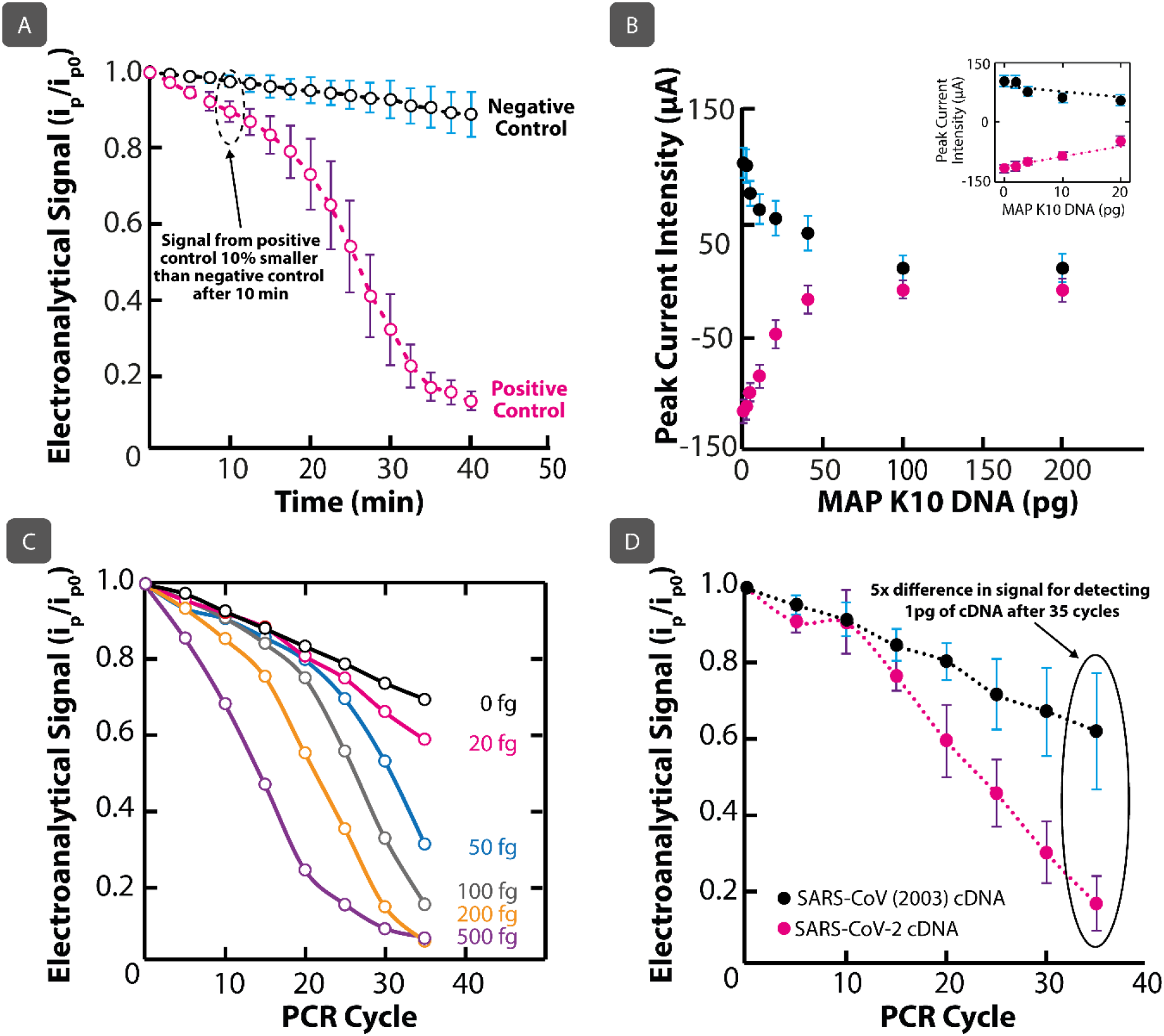
**(A)** qRPA measurements performed using TriSilix. TriSilix produces a clear signal between the positive and negative control sample within 10 min (N=5). **(B)** Titration curve for direct electrochemical detection of genomic DNA from MAP K10 DNA without amplification over 0 to 200 pg (N=3). (**C)** qPCR measurements performed using TriSilix of genomic DNA from MAP K10 against CTMX-ESBL as negative control (0 fg). **(D)** TriSilix qPCR measurement for a synthetic fragment of cDNA from SARS-CoV-2 against a synthetic fragment of cDNA from SARS-CoV (2003) as negative sample. Clear separation of signals after 20 cycles (~40 mins), indicating rapid, on-site detection is possible using TriSilix (N=3). Error bars in all plots=SD.

In the next experiment, we have performed qPCR analysis of genomic DNA extracted from clinical samples of *Mycobacterium avium* subspecies *paratuberculosis* (MAP) K10 strain. MAP is the causative microorganism of Johne’s Disease affecting ruminant animals and is controversially associated with Crohn’s Disease in humans. MAP is, therefore, an important analytical target which requires routine testing in centralized laboratories through microbial cultures or conventional fluorescence qPCR (lasting up to 3 hours). ^29–31^ Using TriSilix, we have first performed a titration experiment to determine the limit-of-detection (LoD) of genomic DNA from MAP K10 without amplification over a range of 0-200 pg (**Figure 5B**). Once again, we performed SWV as the analytical method and measured anodic and cathodic peak current intensities originating from MB as the electroanalytical signal. Without DNA amplification, we were able to detect genomic DNA of MAP K10 (4,829 kbp) down to an LoD of 0.8 pg from cathodic peak current intensities in comparison to a LoD of 1.2 pg provided by the anodic peak current intensities; the cathodic peak current intensity was, therefore, used as the electroanalytical signal in the following PCR experiments. We performed qPCR using the TriSilix chip to amplify and quantify small amounts (0 – 500 fg) of DNA of MAP K10 in real-time (**Figure 5C**). The forward primer (5’-GCC GCG CTG CTG GAG TTG A-3’) and reverse primer (5’- CGC GGC ACG GCT CTT GTT -3’), were used to amplify a 563 nucleotide segment of IS*900* (204–766 of GenBank Accesion Number AE016958.1; National Center for Biotechnology Information, USA), which has 17 repeats in the genome of MAP K10. Using the qPCR approach shown in Figure S7, we were able to measure as low as 20 fg of genomic DNA of MAP K10 which is equivalent to detection of a single bacterium in the sample. We also characterized the same samples using a commercial laboratory qPCR (IDvet, UK) using a test manufactured by ID Gene™ as a gold-standard (**Figure S8**). The results produced by TriSilix were similar or better in comparison to those produced by the commercially available, sophisticated laboratory instrument; for 20 fg of MAP K10 DNA, the resulting C_t_ (cycle threshold) value was 30 using TriSilix, 5 cycles shorter compared to the value obtained using the commercial qPCR (C_t_=35).

In a final experiment, using TriSilix, we performed qPCR analysis of the complimentary DNA (cDNA) of SARS-CoV-2 (22712–22869 nucleotides of GenBank accession number MN908947)^32^ the causative agent of COVID-19, (**Figure 5D**). We have selected the forward primer (5’-CCTA CTA AAT TAA ATG ATC TCT GCT TTA CT-3’) and reverse primer (5’-CAA GCT ATA ACG CAG CCT GTA -3’), to amplify the target sequence. As a negative control, we used a cDNA fragment of another coronavirus, SARS-CoV (17741–17984 nucleotides of GenBank accession number AY274119)^33^ responsible for the SARS outbreak in 2003. Because we did not have access to patient samples, we used cDNA fragments produced synthetically by Integrated DNA Technologies Inc. We used the same PCR thermal cycling and measurement program described in the previous experiment involving MAP K10. Using TriSilix, we were able detect 1 pg of cDNA of SARS-CoV-2 quantitatively, in real-time with specificity against a cDNA sequence from a similar virus (SARS-CoV), in as low as 20 cycles of PCR (lasting ~40 min). After 35 cycles, the difference in the electroanalytical signal was five times, showing clear separation.

## 6. Conclusions

TriSilix is a low-cost, high-precision, integrated nucleic acid amplification and detection technology that is ideally suited for point-of-need diagnosis of infectious pathogens (bacteria, viruses etc.) affecting, humans, animals and plants. TriSilix can perform both isothermal and cyclic nucleic acid amplification; integrates temperature control and DNA detection on the same device, is label-free, requires minimal sample handling and allow operation by minimally trained personnel.

Although TriSilix is a silicon-based technology, it does not need a cleanroom for fabrication; chips can be produced in a standard wet-lab with easily accessible laboratory equipment worldwide. Because fabrication in an advanced silicon foundry is not needed, reliance on the global supply chains is substantially reduced; silicon foundries are located only in a few countries in the world. Each TriSilix chip costs ~0.35 USD (cost of materials is shown in **Table S9**) when produced at wafer-scale with 4-inch wafers but the cost could be reduced further by moving to 8-inch wafers, decreasing the size of each chip and optimizing the fabrication process (for example, the gold contacts at the bottom of the TriSilix chip could be replaced by another appropriate metal to form Ohmic contacts). Due to its small size, integrated form-factor, and low-power requirements, TriSilix can be controlled with a portable, battery-operated handheld analyzer. Using a Li-ion battery with a 4000 mAh capacity (a typical rating for batteries in modern smartphones), TriSilix is estimated to perform at least 40 tests using isothermal amplification reactions at 40 °C lasting 30 min or 13 tests using 35 cycles of PCR (these estimates assume that the electronics for the handheld analyzer also draw 50 mA while operating).

The TriSilix technology has the following three disadvantages: (i) currently TriSilix uses MB as the redox-active reporter to quantify the products of the NA amplification reactions in real-time. MB is known to polymerize and its activity also slightly decreases over time at elevated temperatures. This issue, however, can be addressed by the use of redox-active metal-complexes that also intercalate with DNA, several of which have already been reported in the literature for real-time NA amplification^10,23,34–36^ (ii) TriSilix currently requires purified DNA samples for operation. (iii) Because sample reservoir and polymer films are attached thermally at 110 °C, TriSilix cannot be operated beyond this temperature without causing debonding of the layers and leaks from the sample reservoir. For most NA amplification processes, however, 110 °C is sufficiently high. In case higher temperatures are needed (for example for on-chip sample prep), the fabrication process would need to be modified and thermoplastics with a higher melting point could be used.

In the future, we will design a handheld analyzer and include all electronics needed for the operation of TriSilix in the same device to simplify and enable its use in the field (currently the potentiostat for electrochemical NA sensing and driver electronics for heating + temperature sensing are separate). We will also add capabilities to communicate with smartphones to have access to the cloud so that the result of each diagnostic test can be passed to health agencies remotely.^37,38^ To enable on-site testing, we will implement on-chip lysis capabilities (such as mechanical^39^ or heat-induced cell lysis^40^) to analyze the samples taken directly from the subject without further sample prep. We will also implement on-chip reverse transcription to enable analysis of RNA. TriSilix would be particularly useful in emergency situations, such as the COVID-19 pandemic, where rapid, early, on-site detection of the virus would accelerate intervention and reduce the spread of the pathogen. Since TriSilix is a low-cost, high-performance NA detection technology, it is expected to find applications beyond infectious diseases which may include genetic analysis of various plants, animals or detection of genetic diseases. TriSilix is a versatile platform and could also be adapted for electrochemical detection of non-genetic targets (*e.g.,* through the use of antibodies) with on-chip incubation capabilities to deliver accelerated results at the Point-of-Need.

## 7. Methods and Materials

### Fabrication

#### Preparation of pSi

Silicon wafers (p-type Siegert Wafer, 525 ± 25 μm thickness, 0−100 Ω·cm, lightly doped) were cleaned with acetone, rinsed with distilled water, and then immersed in a 3:1 piranha solution (95% H_2_SO_4_/30% H_2_O_2_ v/ v) at 80 °C to remove organic residues. The wafers were dried in air and immersed in a batch containing an aqueous solution of 80 μM KAuCl_4_ and 0.5% HF for 20 s to deposit Au particles on the surface of the wafer electrolessly. The Si wafers were rinsed with distilled water, dried and immersed in an etching bath (30% H_2_O_2_: 10% HF, 1:20 v/v ratio) for MACE with for 10 min. All chemicals, unless otherwise stated, were purchased from Sigma.

#### Patterning and heat-transfer of polymer layers

PE sheets were laser-cut (Model: Trotec Speedy 100; Wavelength: 1μm; Power: 20 W) and the patterns were transferred to the porous surfaces of the Si wafer (both front and back) by heat-pressing at 180 °C for 5 min. The regions cut out were later electroplated to form the WE and electrical contacts. The Cu-PET (20 μm Cu; 23 μm PET) sheets were purchased from UK Insulations Ltd. and laminated with an additional sheet of PE film at 180 °C for 5 min in a heat press to improve thermal bonding on pSi surfaces. The Cu-PET-PE laminates were also laser-cut using a fiber laser, electroplated with Ag and heat-pressed on the top side of the Si wafer to form the electrodes for the CE and RE.

#### Electrodeposition of Au ang Ag

Immediately before electroplating with Au to form the electrical contacts at the bottom of the wafer and WE on the top, each Si wafer was dipped into 5% HF to remove the native oxide and produce an electrically conductive surface. The wafers were electroplated with Au in a bath containing 10 mM KAu(CN)_2_/KCN using a 5 cm x 5 cm Pt electrode (SPA plating) as anode at a constant current of 10 mA. The Cu-PET-PE film was first treated with 0.1 M H_2_SO_4_, rinsed and electroplated with Ag in an aqueous solution of 10 mM AgCN/KCN under an applied current of 20 mA using a 5 cm x 5 cm Pt electrode as anode. All chemicals, unless otherwise stated, were purchased from Sigma.

### DNA Amplification and Extraction

#### RPA

The RPA Premix solution was prepared by adding and vortexing 25 μL of 2X buffer, 8 μL of 10 mM dNTPs (Fisher Scientific), 5 μL of 10X E Mix and 5 μL of control primer Mix (30 bp) in a 0.2 mL Eppendorf tube. 2.5 μL of 20X core solution and 1 μL of 0.05 % MB were placed in the lid of the tube and mixed with 10 inversions. The RPA Mastermix was made by mixing 3.5 μL of Nuclease-free ultrapure water (Fisher Scientific) with the Premix. This solution was subsequently mixed with 2.5 μL of 280 mM OAc and 1 μL of positive control DNA or 50 nM negative control (CTX-M ESBL: 5’-ATTGACGTGC TTTTCCGCAA TCGGATTATA GTTAACAAGG TCAGATTTTT TGATCTCAAC TCGCTGATTT AACAGATTCG GTTCGCTTTC ACTTTTCTTC-3’; Sigma) to create the final RPA mix. 30 μL of the final solution were added into the device chamber followed by 10 μL of mineral oil (Sigma) to prevent evaporation of water. All of the chemicals (unless otherwise stated) were included in the TwistAmp® Basic Kit.

#### MAP K10 PCR

50 μL of PCR mastermix solution in Nuclease-free ultrapure water contained 60 mM Tris-HCl Buffer pH 8.8 (Bio-Rad), 200 μM dNTPs, 200 nmol μL^−1^ of each primer (Biomers), 0.5 U Taq DNA polymerase and 20 μg mL^−1^ MB. 50 μL of PCR mix solution in nuclease-free water contained 60 mM Tris-HCl Buffer pH 8.8 (Bio-Rad), 200 μM dNTPs, 200 nmol μL^−1^ of each primer (Biomers), 0.5 U Taq DNA polymerase, 2 mM MgCl_2_, 20 μg mL^−1^ MB and 1 μL of MAP or Nuclease-free ultrapure water. 30 μL of PCR mastermix or PCR mix solutions were added into the sample reservoir of TriSilix for reaction, followed by 10 μL of mineral oil (Sigma) to prevent evaporation. All chemicals, unless otherwise stated, were purchased from Fisher Scientific.

#### SARS-CoV PCR

The PCR Mix consisted of 0.5 U Taq polymerase, 1X Taq Polymerase buffer, 1 μM of each primer (Biomers), 2 mM MgCl_2_, 0.2 mM dNTPs, 30 μg mL^−1^ MB and 1 pg cDNA (IDtdna). 30 μL of this solution were added into the sample reservoir of TriSilix for reaction, followed by 10 μL of mineral oil (Sigma) to prevent evaporation. All chemicals, unless otherwise stated, were purchased from Fisher Scientific.

#### Extraction of Genomic DNA

Genomic DNA was prepared from a 10ml culture of MAP K10 in Middlebrook 7H9 supplemented with 10% Albumin Dextrose Catalase enrichment, 0.2% (vol/vol) glycerol, 0.05% (vol/vol) Tween 80 and 2μg/ml mycobactin j. Mycobacteria were harvested in early to mid log phase of growth and the cells were pelleted at room temperature for 5 min at 14,000g. Pellets were re-suspended in ATL buffer (Qiagen DNeasy® Blood & Tissue Kit) and the samples were transferred to Lysing Matrix B tubes (0.1mm silica spheres). Samples were homogenized in a FastPrep™ FP120 cell disruptor at 3 x 20s, Speed 6, followed by centrifugation at 14,000g for 5 minutes. Supernatants were transferred to fresh sterile micro centrifuge tubes and Proteinase K (Qiagen DNeasy® Kit) was added followed by incubation overnight at 56°C. DNA was extracted using DNeasy® Kit (Qiagen) according to the manufacturer’s protocol. Concentration (40 μg mL^−1^) and purity of the DNA were determined using a Nanodrop and the absorbance ratios 260/280 and 260/230.

### Characterization

#### Microstructural Characterization

Scanning electron micrographs were acquired using a Zeiss Gemini Sigma 300 FEG SEM at 5 keV electron beam energy. Optical micrographs were obtained with a Brunel SP202XM metallurgical microscope. A Nikon D3200 camera was attached to the microscope for the acquisition of the images using DigiCamControl open-source software.

#### Electrical and Thermal characterization

The I-V curves for the characterization of the metal−semiconductor junctions were acquired using a Keithley 2450 SourceMetter. Thermal images were recorded using a FLIR E4 thermal camera.

#### Electrochemical Measurements

Cyclic (CV) and square wave (SWV) voltammograms were acquired using a handheld potentiostat (PalmSens3, PalmSens BV, The Netherlands) with the supplied PSTrace 5.3 software in a three-electrode setup. The following parameters were used for all the SWV measurements: pulse amplitude of 100 mV, potential step of 5 mV and 50 Hz of frequency. Prior to the DNA measurements by SWV, the sample reservoir was cleaned with DNA decontamination reagent (Fisher Scientific) followed by Nuclease-free ultrapure water (Fisher Scientific).

## 8. Acknowledgements

F.G., K.S. and E.N-B. would like to thank the Wellcome Trust (Grant No. 207687/Z/17/Z) and the UK Engineering and Physical Sciences Research Council (EPSRC) (EP/ R010242/1) and Imperial College, Department of Bioengineering for their generous support. M. Kasimatis acknowledges EPSRC DTP (Reference: 1846144) and A.C., BBSRC DTP (Reference: 2177734). Y.C. thanks the Turkish Ministry of Education. M. Kaisti acknowledges Finnish Foundation for Technology Promotion. F.G. and M.G. thank the Imperial College Centre for Plastic Electronics. F.G. also acknowledges Agri Futures Lab. We would like to thank George Caldow from SRUC (Midlothian, UK) for performing the commercial qPCR MAP tests and Dr Tolga Bozkurt (Imperial College) for the fruitful discussions.

