## Supplemental Information for "Ultra-Low-Cost Integrated Silicon-based Transducer for On-Site, Genetic Detection of Pathogens"

**Video V1.** Animation about the components and performance of TriSilix (Tri-modal Silicon-based integrated nucleic acid transducer). The video starts showing the dimensions and components of the chip to perform qPCR: sample reservoir, PCR reagents with redox reporter and silicon-based transducer. Then, it explains the working principles and signaling of the three modes of operation of the silicon-based transducer: i) electrochemical qPCR; ii) electrical heating (Joule heating) and, iii) thermal sensing (thermistor).

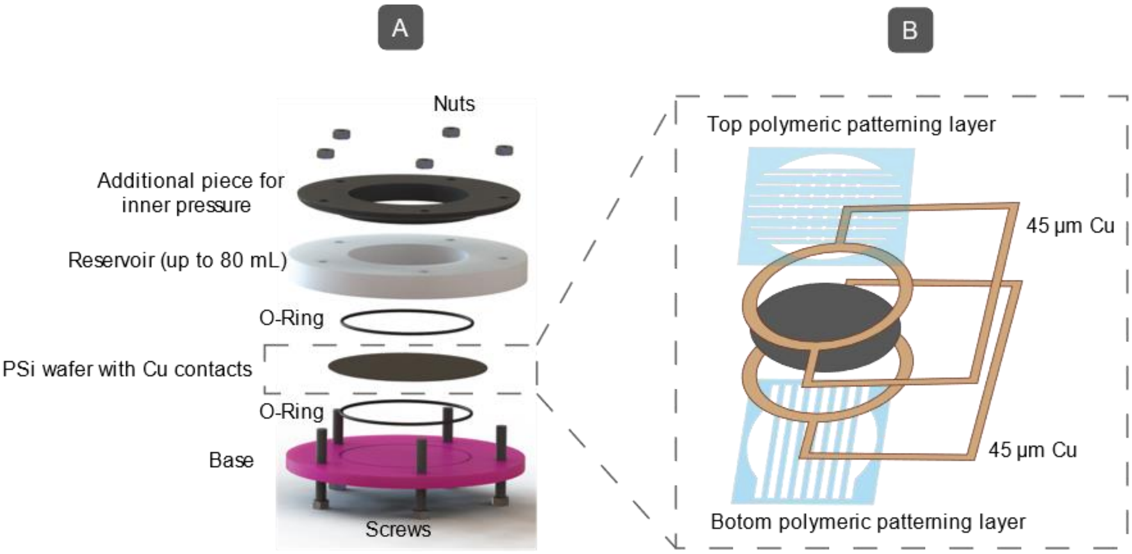


**Figure S1 – Custom holder for electroplating.** Schematic illustrations of **A)** the holder and **B)** integration of thin copper contacts for metal electroplating.

To achieve high uniformity when electroplating across the wafer, we have designed, and 3D printed with polylactic acid (PLA) a custom holder which has a circular contact around the edges of the wafer (Figure S1A). After the incorporation of the polymeric patterning layer on the porous silicon wafer, the surface is not completely flat and the rounded copper contacts (40 µm thicker) must be incorporated, at the top and bottom, between the polymeric layer and the silicon wafer in order to guarantee a good electrical contact for the metal electroplating (Figure S1B). Then, the reservoir is placed on the top and the holder, closed through five screws and filled with 20 mL of plating solution. Since we observed a progressive leaking of the solution with the time, rubber O-rings were integrated in both, base and reservoir, pieces and a third component was incorporated in order to apply an inner pressure in the region of the O-rings. Once one of the sides of the wafer is electroplated, the plating solution is extracted, and the holder is opened to flip the wafer with contacts. Finally, the holder is closed again through the screws and the plating solution is added again.

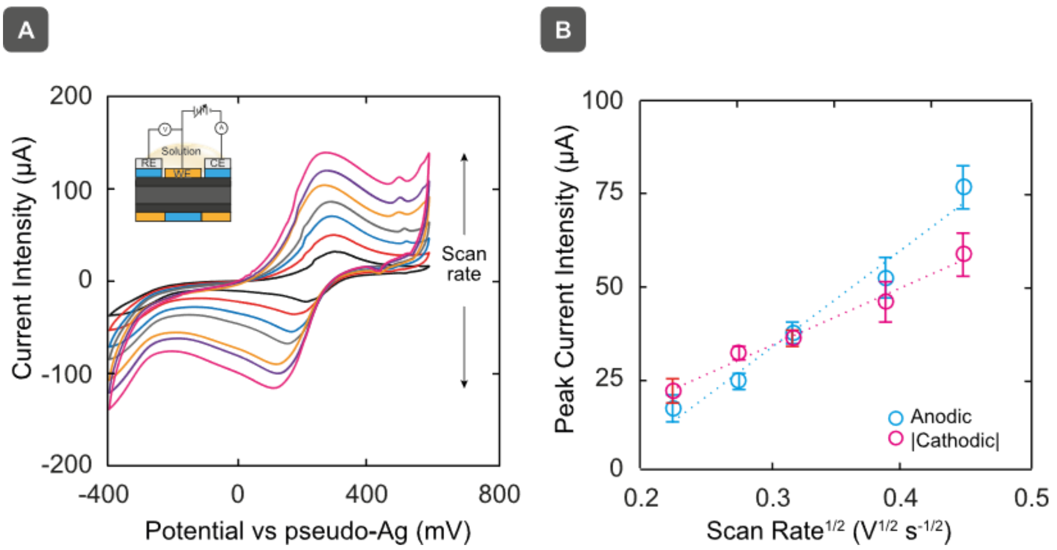


**Figure S2 – Calculation of electrocatalytically active area through Randles-Sevcick. A)** CVs in 2 mM $K_4Fe(CN)_6$ solution (0.1 M KCl) sweeping the potential from -600 to +700 mV at 10, 50, 75, 100, 150 and 200 mV $s^{-1}$. **B)** Peak current intensity vs scan rate plots from CVs recorded using 5 devices.

The electroactive area was evaluated by cyclic voltammetry adding 50 µL of 2 mM $K_4Fe(CN)_6$ solution in 0.1 M KCl on the device. The dependence of the peak current on the scan rate was evaluated by cyclic voltammetry sweeping the potential from -400 to +600 mV vs Ag at 10, 50, 75, 100, 150 and 200 mV $s^{-1}$. The electrochemical cell of the device (Au-PSi WE, Ag CE and RE electrodes) was connected to the potentiostat (PalmSens3 model from PalmSens, UK) with flat crocodile clamps. All reagents were purchased from Sigma.

According to the Randles-Sevcick equation for a flat electrode and for diffusion-controlled processes at 25 °C. [1-4] $\boldsymbol{ip}=(2.69\cdot10^5\ )n^{3/2}\ \boldsymbol{A}\ D^{1/2}\ C^*\ \boldsymbol{v}^{1/2}$. Where $\boldsymbol{ip}$ is the peak current (A), $n$ is the number of electrons transferred (n = 1 for ferrocyanide), $\boldsymbol{A}$ the effective area of the electrode ($cm^2$), $D$ is the diffusion coefficient of ferrocyanide in aqueous solutions ($6.50x10^{-6}$ $cm^2$ $s^{-1}$), $C^*$ is the concentration ($2x10^{-6}$ mol $cm^{-3}$) and $\boldsymbol{v}$ is the scan rate (V $s^{-1}$). Cyclic voltammograms, as those shown in Figure S3a, were recorded using five different devices without washings between scans. The gradient of the logarithmic plot peak current intensity vs the scan rate was 0.71±0.03 ($R^2$ = 0.995) and 0.69±0.05 ($R^2$ = 0.997), for anodic and cathodic processes, respectively. Readjusting the Randles-Sevcik equation with the experimental gradient values, calculated effective areas were 20±0.3 $mm^2$ and 19±0.5 $mm^2$

from anodic and cathodic data, respectively. These values are at least the double of the geometric area (9.6 $mm^2$).

**Section S3 – Calibration of TriSilix Thermistor**

The degree of precision on the temperature control of the NA amplification chamber is related to the accuracy in establishing the resistance–temperature correlation curve for an Ohmic resistor: $R = R_0(1 + \alpha(T-T_0))$ where R and $R_0$ are resistances of the thermistor at temperatures T and $T_0$ (reference temperature), and α is the temperature coefficient of resistance. This equation is used to convert the resistance reading (calculated from the voltage through the diode connected to the custom board, measured at a constant current of 10 mA every 0.5 s) to the temperature reading of the thermal camera. When $R/R_0$ is plotted against $T-T_0$, linear graphs (Figure 3B) with a regression coefficient of 0.9991±0.0007 are obtained using 5 different devices. The slope of this linear graph gives an α value of 6.1±0.4 x 10-3 $°C^{-1}$. With the well-calibrated temperature sensors, fast thermal cycling (heating and cooling gradients of 3.2 and 2.5 $°C{\cdot}s^{-1}$, respectively) of the NA amplification chamber (Figure 3E) is achieved with a temperature precision of ±1.3 °C when the values measured by the thermal camera are compared with those of the recorded resistances by the MATLAB-based interface.

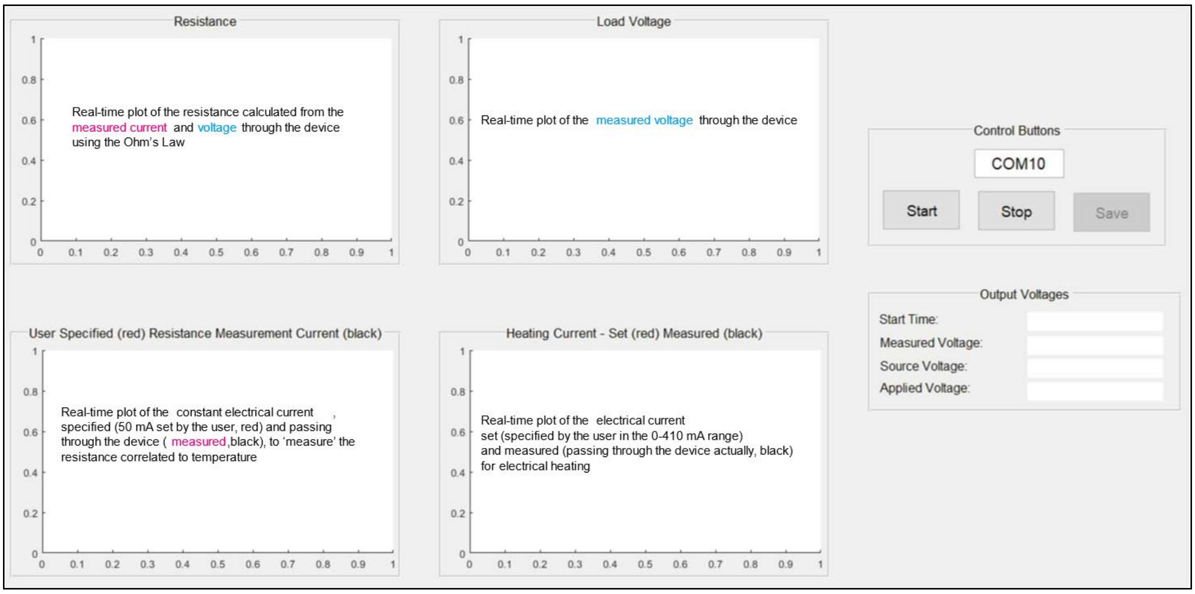


**Figure S4. MATLAB-based graphical user interface.** The X axis of every of the four screens shows the time of the recording in seconds and Y axis shows the resistance in Ohms (upper left screen), the voltage in volts (upper right screen) and the current in amps (bottom screens).

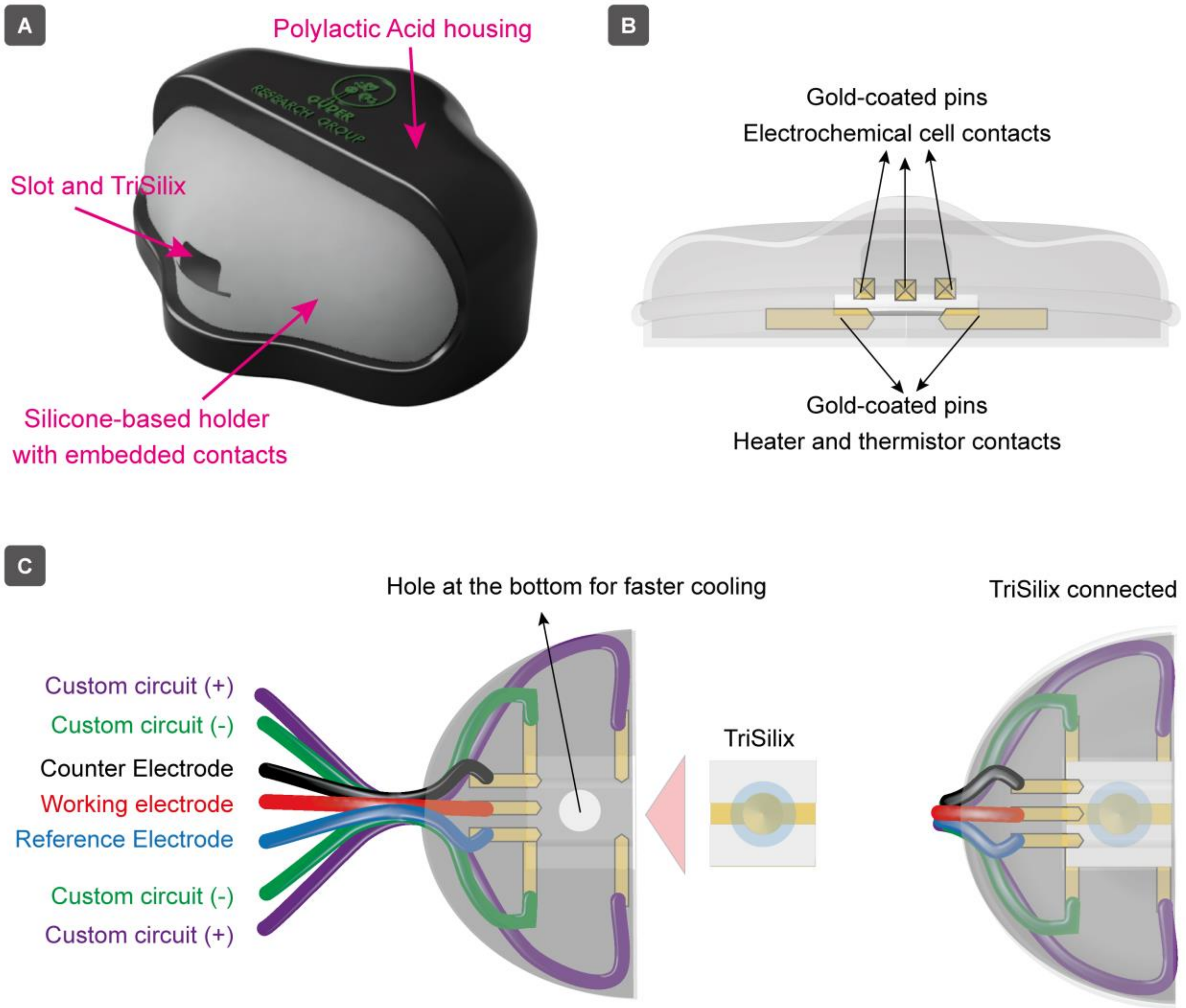


**Figure S5 – Silicone-based sample holder in 3D-printed PLA housing. A)** Front view of the holder with contacts for trimodal performance of Trisilix. Schematic representations of **B)** the distribution of embedded contacts and **C)** connection of Trisilix (top view) for trimodal performance using the holder.

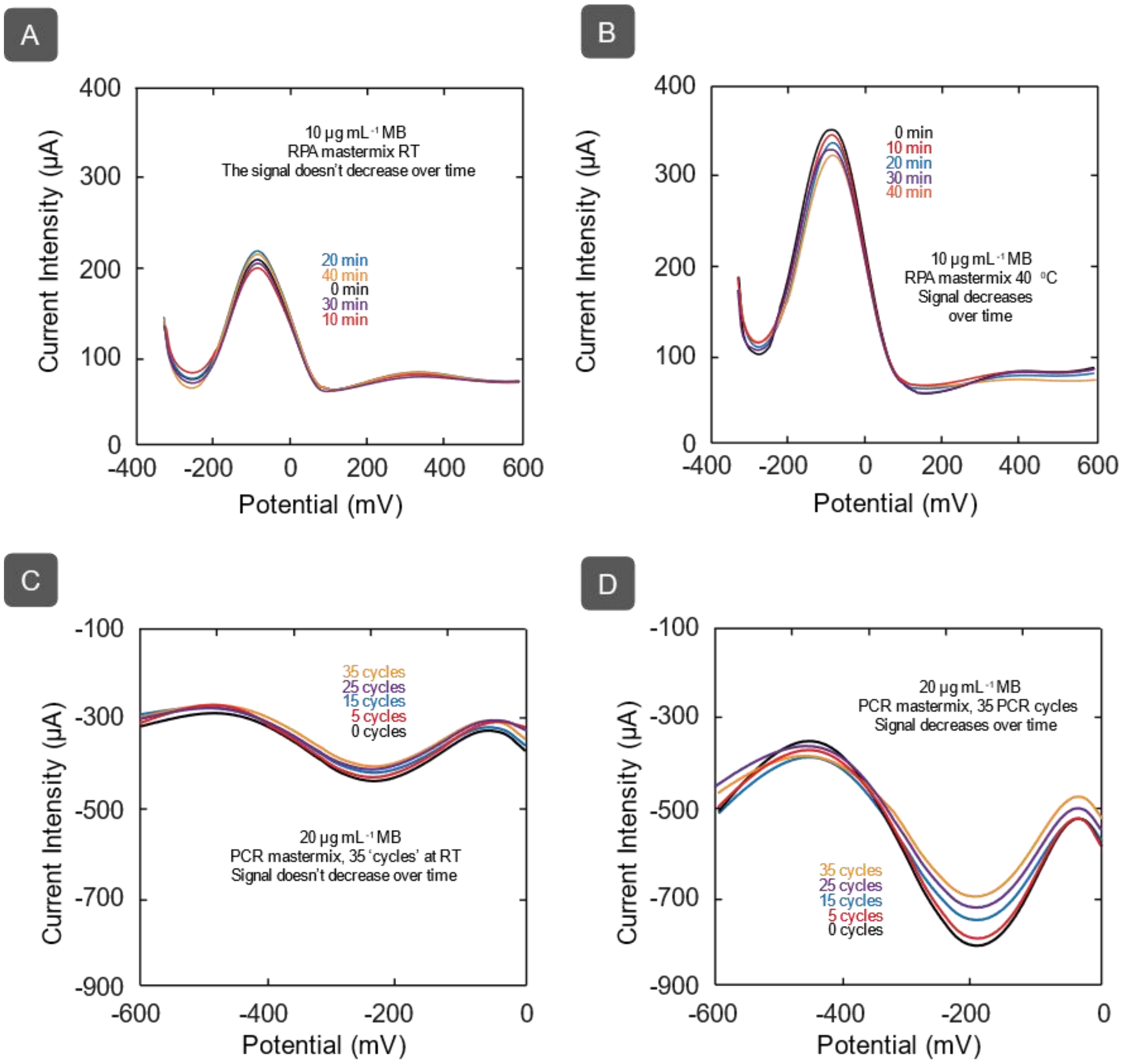


**Figure S6:** Representative SWVs from those recorded every 2.5 min in RPA solutions without DNA (RPA mastermix) under isothermal conditions at **A)** room temperature and **B)** 40°C during 40 min. Representative SWVs from those recorded every 5 cycles in PCR solutions without DNA (PCR mastermix) **A)** at room temperature and **B)** under the programmed thermal cycling conditions during 35 cycles. Esw: 100 mV; Es: 5 mV; f:50 Hz.

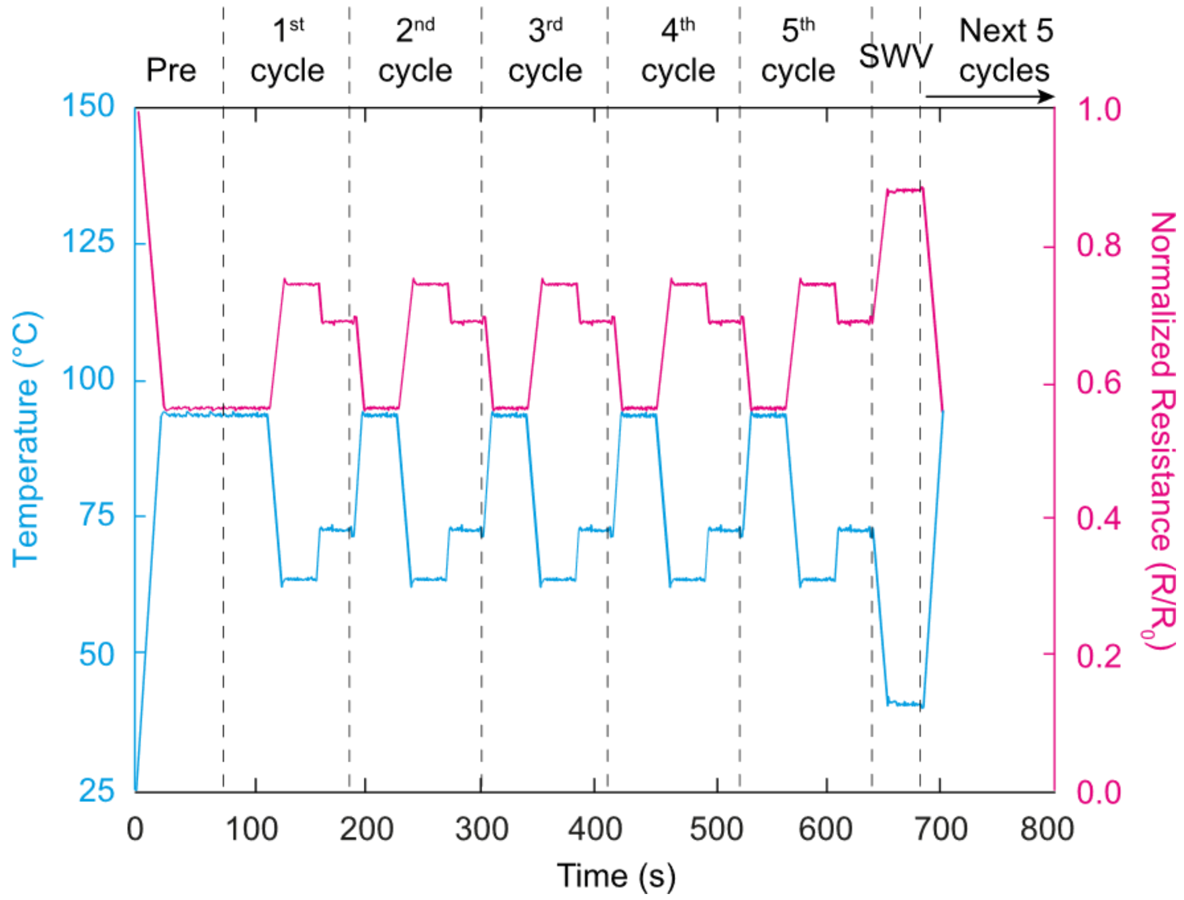


**Figure S7 – Program of temperatures over time during PCR experiments.** Thermal cycling curves of the first 5 cycles in qPCR experiments. Dashed lines delimitate each of the 5 PCR cycles performed before SWV measurements.

Each cycle of qPCR consisted of three steps: (i) denaturing at 94°C for 30 sec, (ii) annealing at 63°C for 30 sec and (iii) extension 72°C for 30 sec. In addition to these three steps, every fifth cycle, we performed SWV at 40°C (maintained for 30 s) to determine the peak current intensity in relation to the redox processes involving MB. The program of currents to heat the chamber during a PCR cycle consisted in a first step of preheating applying a current of 410 mA for 23 s and 400 mA for 30 s. Then, 5 times the following program: i) denaturalization applying 410 mA for 17 s (except for the first cycle) and 400 mA for 30 s; ii) primer annealing applying 0 mA for 10 s and 270 mA for 30 s; iii) extension applying 315 mA for 3 s and 310 mA for 30 s; iv) electroanalysis applying 0 mA for 12 s and 170 mA for 30 s. Finally, to reach 94 °C for the start of the next series of 5 cycles, a current of 410 mA is applied during 17s.

-

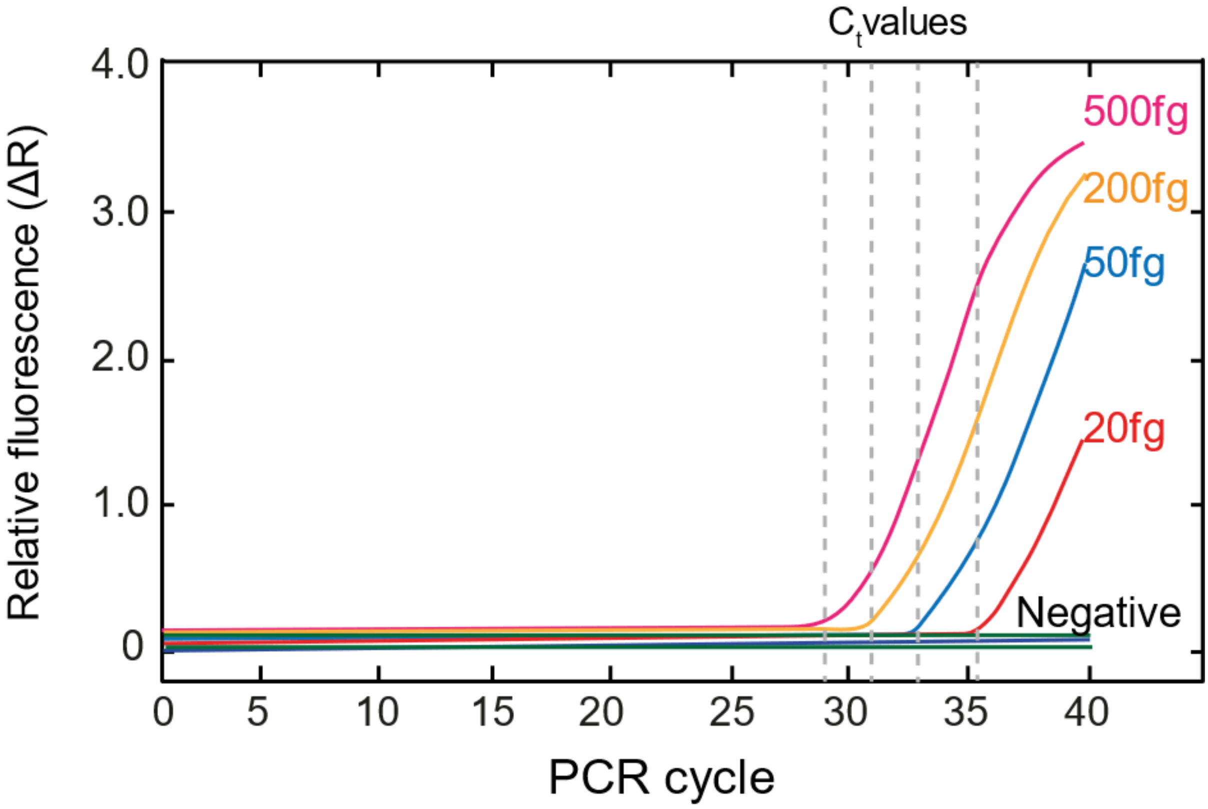


**Figure S8 – qPCR results of MAP K10 DNA using a standard method.** MAP K10 DNA amplification plot using the IDVet commercial qPCR test for Johne's disease. The set cut-off for positives is $C_t$ 40 and samples with Ct below 33 are re-tested as standard to confirm the original positive result.

**Table S9 –Table showing the breakdown of the cost after wafer-scale manufacturing**

| Product | Conc | Supplier | Reference | Amount | Purchase Price | Spent Quantity | Spent Cost | Number of batches (37 devices) | Cost per device |
|---|---|---|---|---|---|---|---|---|---|
| **HF** | 50% | Sigma | 30107-500ML-M | 500 mL | 3.6 | 13.8 mL | 0.1 | 1 | **0.0027** |
| **$H_2O_2$** | 30% | VWR | 8.22287.2500 | 2500 mL | 16.57 | 13 mL | 0.09 | 1 | **0.0023** |
| **$H_2SO_4$** | 98% | VWR | 20700.42 | 2500 mL | 9.43 | 30 mL | 0.11 | 1 | **0.0031** |
| **$KAuCl_4$** | 100.00% | Sigma | 450235-250MG | 250 mg | 62.19 | 300 mg | 0.75 | 1 | **0.0202** |
| **$KAu(CN)_2$** | 99.95% | Sigma | 379867-250MG | 250 mg | 108.68 | 116 mg | 5.01 | 3 | **0.0451** |
| **KCN** | ≥98% | Sigma | 60178-25G | 25 g | 19 | 52 mg | 0.04 | 3 | **0.0004** |
| **AgCN** | 99 % | Sigma | 184535-10G | 10 g | 48.68 | 73 mg | 0.36 | 3 | **0.0032** |
| **Cu-PET** | 20/23 um | UK Ins. | | 650mmx25m (162,500 $cm^2$) | 3.53 | 5 pieces 256 $cm^2$ | 0.03 | 1 | **0.0008** |
| **PE** | | Kite UK | | 114x114mm 100 units | 2.71 | 3 units | 0.08 | 1 | **0.0022** |
| **Si wafer** | | Inseto | | 1 unit | 11.71 | 1 unit | 9.92 | 1 | **0.2681** |
| | | | | | | | | TOTAL | 0.35 |

* All prices are in USD
